# Glomerular function and urinary biomarker changes between vancomycin and vancomycin plus piperacillin-tazobactam in a translational rat model

**DOI:** 10.1101/2021.11.08.467852

**Authors:** Jack Chang, Gwendolyn Pais, Kimberly Valdez, Sylwia Marianski, Erin F. Barreto, Marc H. Scheetz

## Abstract

Clinical studies have reported additive nephrotoxicity associated with the combination of vancomycin (VAN) and piperacillin-tazobactam (TZP). This study assessed differences in glomerular filtration rate (GFR) and urinary biomarkers between rats receiving VAN and those receiving VAN+TZP. Male Sprague-Dawley rats (n=26) were randomized to receive 96 hours of intravenous VAN at 150mg/kg/day, intraperitoneal TZP at 1400 mg/kg/day, or VAN+TZP. Kidney function was evaluated using fluorescein-isothiocyanate sinistrin and a transdermal sensor to estimate real-time glomerular filtration rate (GFR). Kidney injury was evaluated via urinary biomarkers including kidney injury molecule-1 (KIM-1), clusterin, and osteopontin. Compared to a saline control, only rats in the VAN group showed significant declines in GFR by day 4 (−0.39 mL/min/100 g body weight, 95% CI: -0.68 to -0.10, p=0.008). When the VAN+TZP and VAN alone treatment groups were compared, significantly higher urinary KIM-1 was observed in the VAN alone group on day 1 (18.4 ng, 95% CI: 1.4 to 35.3, p=0.03), day 2 (27.4 ng, 95% CI: 10.4 to 44.3, p=0.002), day 3 (18.8 ng, 95% CI: 1.9 to 35.8, p=0.03), and day 4 (23.2 ng, 95% CI: 6.3 to 40.2, p=0.007). KIM-1 was the urinary biomarker that most correlated with decreasing GFR on day 3 (Spearman’s rho: -0.45, p = 0.022) and day 4 (Spearman’s rho: - 0.41, p = 0.036). Kidney function decline and increased KIM-1 were observed among rats that received VAN only, but not TZP or VAN+TZP. Addition of TZP to VAN does not worsen kidney function or injury in a validated translational rat model.

## Introduction

Vancomycin (VAN) is a glycopeptide antibiotic that is a treatment of choice for many resistant Gram-positive infections. It remains the most frequently prescribed parenteral antibiotic in United States hospitals and use has steadily increased over the past decade (1). Nephrotoxicity is a common adverse effect of VAN, with attributed rates of >10% (2-4). Piperacillin-tazobactam (TZP) is a beta-lactam/beta-lactamase inhibitor combination that is commonly co-prescribed with vancomycin for empiric coverage of Gram-negative organisms (5, 6).

Recent clinical meta-analyses have found the combination of VAN and TZP to be associated with additive nephrotoxicity, as assessed by increased serum creatinine (SCr), compared to VAN alone (7-11). Luther et al. found that the combination of VAN+TZP increased the odds of acute kidney injury (AKI) when compared to VAN monotherapy (OR: 3.4, 95% CI: 2.57-4.50) or TZP monotherapy (OR: 2.7, 95% CI: 1.97-3.69). This difference in AKI also persisted when VAN+TZP was compared to VAN+cefepime or meropenem (OR: 2.68, 95% CI: 1.83-3.91) (9). More recently, Bellos et al. recapitulated these findings in the largest meta-analysis of VAN+TZP nephrotoxicity to date. Their study included over 50,000 patients and VAN+TZP was found to increase the odds of AKI, as assessed by SCr using the Kidney Disease: Improving Global Outcomes guideline, when compared to VAN (OR: 2.05, 95% CI: 1.17-3.46), VAN+cefepime (OR: 1.80, 95% CI: 1.13-2.77), and VAN+meropenem (OR: 1.84, 95% CI: 1.02-3.10) (10). Despite these results from retrospective studies suggesting that VAN+TZP is associated with greater increases in SCr compared to VAN alone, it is still unclear whether this reflects overt nephrotoxicity. The underlying mechanism of additive nephrotoxicity associated with VAN+TZP remains unknown and SCr is known to be a poor surrogate marker for both kidney function and damage (12).

Clinical study outcomes remain limited by mostly retrospective research design and use of SCr as the sole indicator of kidney injury and function. However, SCr is not a direct marker of kidney injury; instead, it is an imperfect surrogate for glomerular filtration rate (GFR) (12-14) and changes in the secretion and/or reabsorption of creatinine can result in fluctuations in SCr that do not reflect true loss of kidney function. An additional limitation of SCr is that during acute changes in kidney function, GFR can decline by as much as 50% before detectable rises in SCr occur (14). To address the limitations of using SCr as a surrogate for kidney function and injury, several translational animal studies have investigated the injury profile of VAN+TZP vs. VAN using newer kidney biomarkers of injury (e.g. kidney injury molecule-1 [KIM-1], clusterin, and osteopontin (OPN)) and histopathology (15). In these models, results conflicted with retrospective human studies; additive nephrotoxicity associated with VAN+TZP was not observed in animal studies (15, 16). KIM-1 was identified as the most relevant urinary biomarker in animal studies for vancomycin induced kidney injury, correlating with GFR changes and end-organ histopathologic damage at proximal tubule (15, 17-19). While injury biomarkers and histopathology scores were not increased in rats receiving VAN+TZP vs. VAN, the impact of TZP on glomerular function was not studied directly.

Thus, the purpose of this study was to employ our translational rat model to directly assess kidney function differences between treatment groups by GFR, and to correlate GFR to urinary injury biomarker expression in rats receiving VAN +/-TZP, TZP alone, or control (saline).

## Results

### Characteristics of animal cohort

A total of 26 male Sprague Dawley rats were studied, with the animal dosing group assignment shown in Table 1. One animal provided only terminal plasma samples due to occluded catheters; all other animals contributed complete data. One animal had missing GFR measurements on one experimental day due to sensor malfunction; all other animals contributed complete GFR data to the model. Mean weight change was not different between the VAN, TZP, VAN+TZP, and saline control groups (−1.39 g vs -2.27 g vs -1.59 g vs +5.50 g, p = 0.19).

**Table 1:** Marginal differences versus saline as referent group.

| Dosing day | Treatment groups |  |  |
| --- | --- | --- | --- |
|  | VAN | TZP | VAN+TZP |
| Urine output difference (mL), mean (95% CI), p-value |  |  |  |
| Baseline | 0.63 (-3.19 to 4.45), p=0.747 | 2.85 (-1.12 to 6.82), p=0.159 | 3.33 (-0.49 to 7.15), p=0.088 |
| Day 1 | 4.09 (0.26 to 7.91), **p=0.036** | 1.40 (-2.57 to 5.37), p=0.490 | 5.52 (1.69 to 9.34), **p=0.005** |
| Day 2 | 0.45 (-3.38 to 4.27), p=0.820 | -1.98 (-5.95 to 1.99),<br>p=0.328 | 0.75 (-3.08 to 4.52), p=0.703 |
| Day 3 | 0.49 (-3.33 to 4.32), p=0.801 | 0.58 (-3.39 to 4.55), p=0.773 | 3.19 (-0.63 to 7.02), p=0.102 |
| Day 4 | 8.87 (5.05 to 12.7), **p<0.001** | -0.07 (-4.04 to 3.91),<br>p=0.974 | 6.83 (3.01 to 10.7), **p<0.001** |
| Urinary KIM-1 difference (ng), mean (95% CI), p-value |  |  |  |
| Baseline | -1.24 (-18.9 to 16.4), p=0.891 | -0.45 (-18.8 to 17.9),<br>p=0.961 | 1.22 (-16.4 to 18.9), p=0.892 |
| Day 1 | 21.5 (3.83 to 39.2), **p=0.017** | -1.42 (-19.8 to 16.9),<br>p=0.879 | 3.14 (-14.5 to 20.8), p=0.728 |
| Day 2 | 29.8 (12.2 to 47.5), **p=0.001** | -3.74 (-22.1 to 14.6),<br>p=0.689 | 2.47 (-15.2 to 20.1), p=0.784 |
| Day 3 | 21.6 (3.91 to 39.2), **p=0.017** | -2.66 (-20.9 to 15.7),<br>p=0.776 | 2.74 (-14.9 to 20.4), p=0.761 |
| Day 4 | 24.1 (6.41 to 41.7), **p=0.008** | -4.14 (-22.5 to 14.2),<br>p=0.658 | 0.83 (-16.8 to 18.5), p=0.927 |
| Urinary clusterin difference (ng), mean (95% CI), p-value |  |  |  |
| Baseline | 108 (-1835 to 2052), p=0.913 | 507 (-1511 to 2524),<br>p=0.623 | 1783 (-160 to 3727), p=0.072 |
| Day 1 | 1563 (-381 to 3507), p=0.115 | 930 (-1087 to 2947),<br>p=0.366 | 2870 (926 to 4814), **p=0.004** |
| Day 2 | 2030 (86 to 3974), **p=0.041** | -597 (-2614 to 1420),<br>p=0.562 | 883 (-1061 to 2826), p=0.374 |
| Day 3 | 1760 (-184 to 3704), p=0.076 | 43 (-1975 to 2060), p=0.967 | 1843 (-101 to 3787), p=0.063 |
| Day 4 | 523 (-1421 to 2467), p=0.598 | -228 (-2246 to 1789),<br>p=0.824 | 400 (-1544 to 2344), p=0.687 |
| Urinary OPN difference (ng), mean (95% CI), p-value |  |  |  |
| Baseline | 0.21 (-0.55 to 0.98), p=0.586 | 0.17 (-0.62 to 0.97), p=0.670 | 0.15 (-0.61 to 0.92), p=0.695 |
| Day 1 | 0.47 (-0.29 to 1.23), p=0.229 | 0.54 (-0.26 to 1.33), p=0.186 | 0.47 (-0.29 to 1.23), p=0.233 |
| Day 2 | 1.09 (0.33 to 1.86), **p=0.005** | 0.35 (-0.44 to 1.15), p=0.383 | 0.51 (-0.25 to 1.28), p=0.189 |
| Day 3 | 1.35 (0.58 to 2.11), **p=0.001** | 0.71 (-0.09 to 1.49), p=0.082 | 0.89 (0.13 to 1.66), **p=0.022** |
| Day 4 | 1.73 (0.97 to 2.49), **p<0.001** | 0.49 (-0.29 to 1.29), p=0.222 | 1.77 (1.01 to 2.53), **p<0.001** |
| GFR difference (mL/min/100 g B.W.), mean (95% CI), p-value |  |  |  |
| Baseline | 0.18 (-0.11 to 0.47), p=0.234 | 0.18 (-0.12 to 0.48), p=0.232 | 0.05 (-0.23 to 0.34), p=0.712 |
| Day 1 | -0.16 (-0.46 to 0.14), p=0.285 | 0.02 (-0.29 to 0.32), p=0.922 | -0.16 (-0.44 to 0.13), p=0.293 |
| Day 2 | -0.11 (-0.39 to 0.18), p=0.474 | 0.0 (-0.30 to 0.30), p=0.999 | 0.01 (-0.28 to 0.31), p=0.938 |
| Day 3 | -0.01 (-0.30 to 0.28), p=0.934 | 0.15 (-0.15 to 0.45), p=0.339 | 0.17 (-0.12 to 0.46), p=0.256 |
| Day 4 | -0.39 (-0.68 to -0.10), **p=0.008** | -0.13 (-0.43 to 0.17),<br>p=0.402 | -0.04 (-0.33 to 0.25), p=0.794 |

**Table 2:**
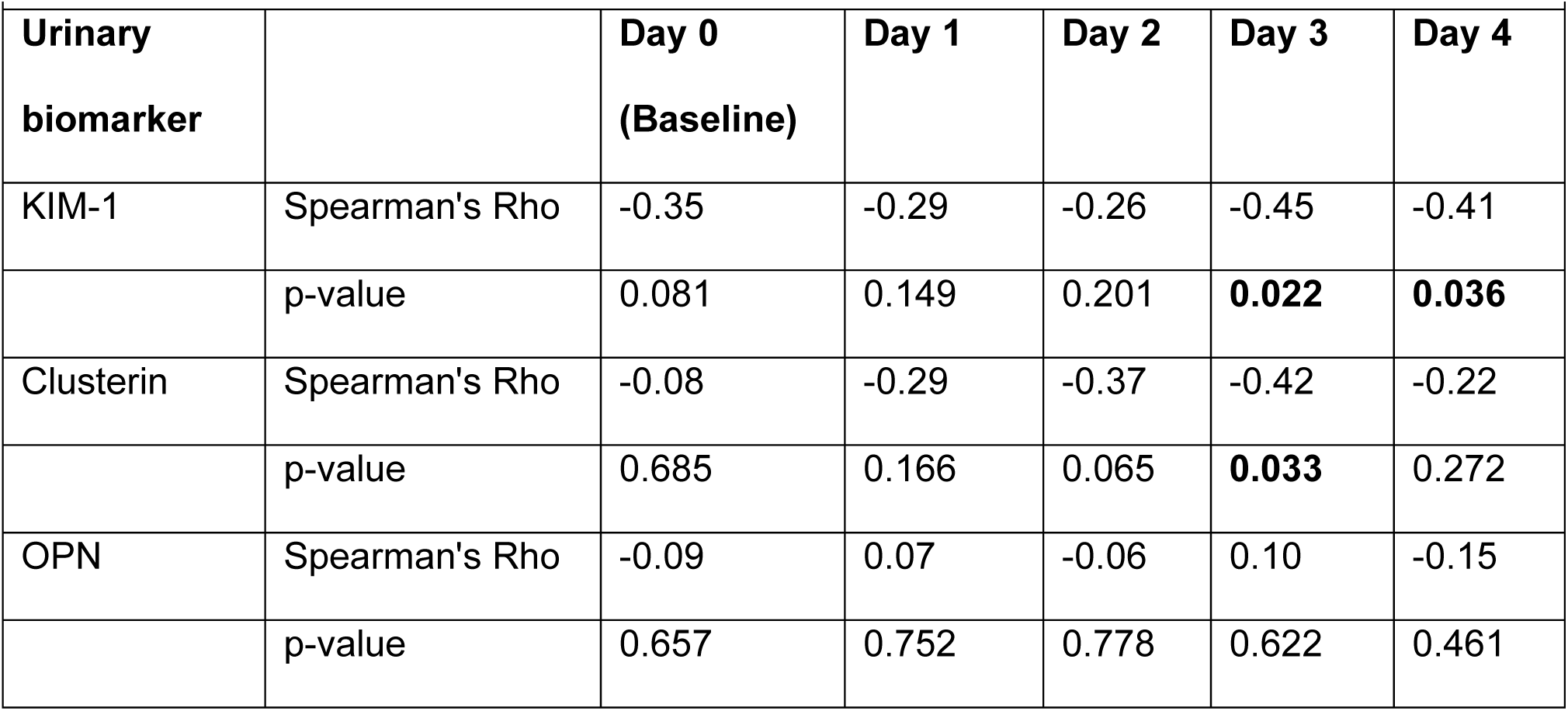
Summary of urinary biomarker correlations with GFR.

### GFR over time

Baseline GFR was not different between the VAN, TZP, VAN+TZP, and saline control groups (with saline used as the referent group) [1.08 ± 0.27, 1.09 ± 0.38, 0.96 ± 0.21, 0.91 ± 0.53 mL/min/100 g body weight, p=0.756]. Following administration of the first dose (day 1), all rats experienced a non-significant decline in GFR. Rats in the VAN group did not recover the GFR decrease and had progressive functional decline, whereas rats that received VAN+TZP, TZP, or saline control subsequently recovered their GFR over the remaining dosing days (Figure 1). When compared to the saline control group, only rats which received VAN had a significant decline in GFR by day 4 (Figure 2, -0.39 mL/min/100 g body weight, 95% CI: -0.68 to -0.10, p=0.008). In a direct comparison of the VAN+TZP and VAN alone groups, the same trend was observed; only rats which received VAN had a significant decline in GFR by day 4 (Supplemental figure 4, -0.35 mL/min/100 g body weight, 95% CI: -0.63 to -0.07, p=0.013).

**Figure 1:**
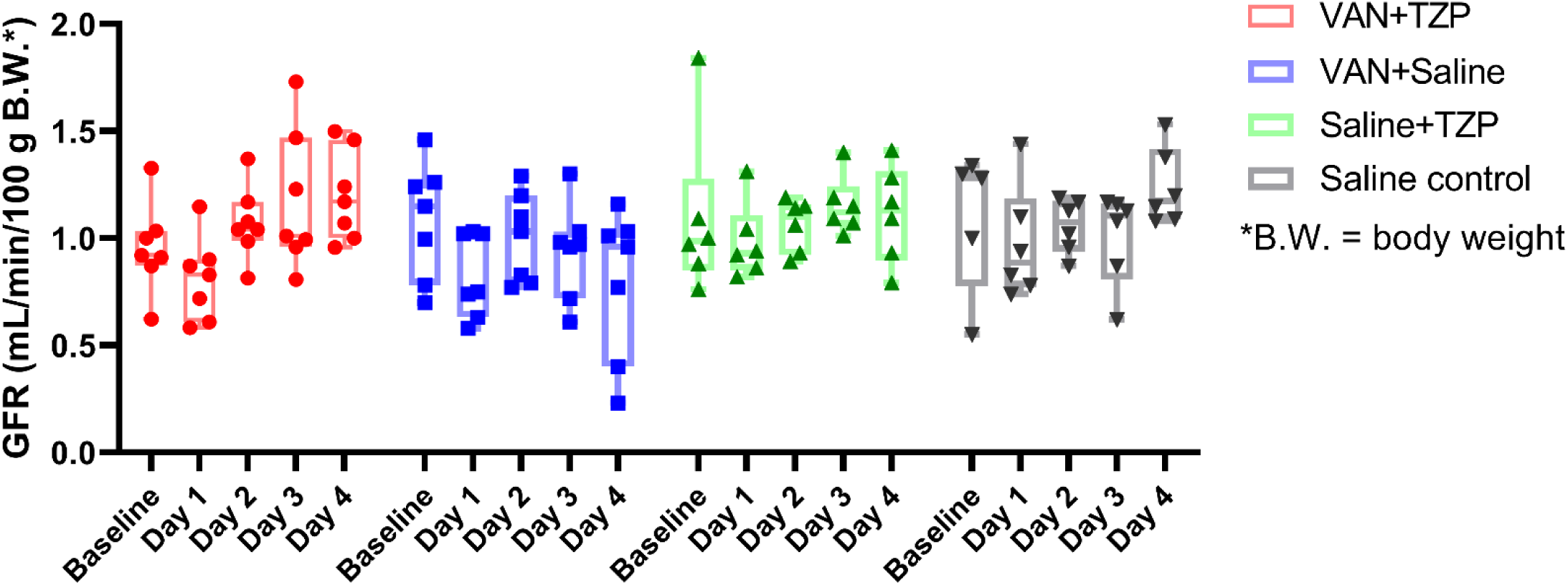
GFR comparison between groups across dosing days. Comparison of individual GFR measurements between treatment groups, and across dosing days. The majority of the rats experienced a decline in GFR after the first dose of study treatment was given on day 1. Rats in the VAN+saline (Blue) group showed progressive functional decline over the study days while rats in all other groups recovered their GFR to baseline levels by day 4. Individual rats are depicted by each data point.

**Figure 2:**
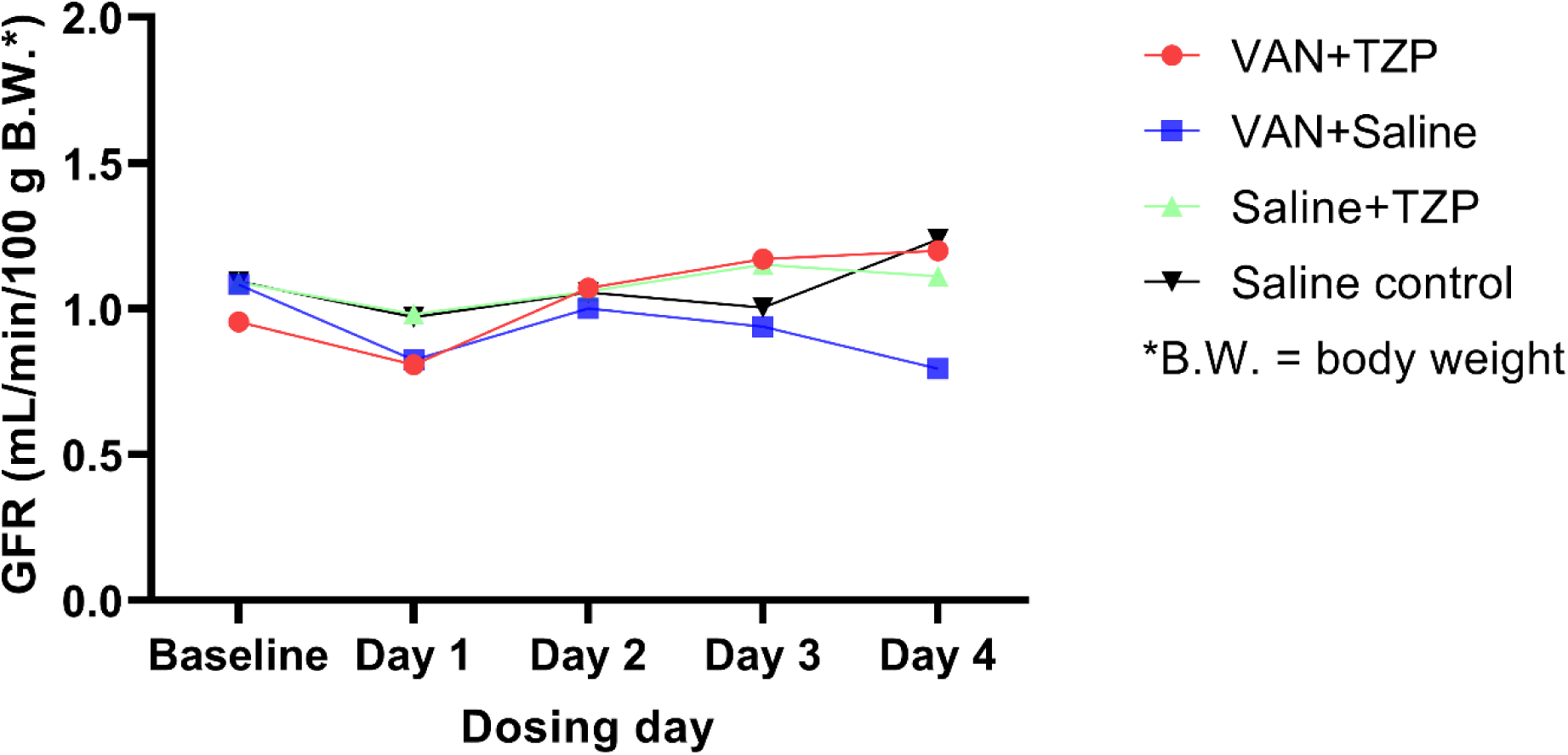
Mean GFR comparison among groups. Comparison of mean GFR measurements between treatment groups, and across dosing days. When the saline control was used as the referent group, only rats which received VAN had a significant decline in GFR by day 4 (−0.39 mL/min/100 g body weight, 95% CI: -0.68 to -0.10, p=0.008). Daily group mean GFR measurements are depicted by each data point.

### Urine output and injury biomarkers

Summary statistics for urine output and urinary biomarker data are listed in Table 1. Baseline urine output was not different among the treatment groups. Daily urine output was compared to baseline and significant differences were seen on day 1 in rats that received VAN (mean difference: -6.9 mL, 95% CI: -11.7 to -2.2, p=0.009) and day 1 in the saline control group (mean difference: -3.5 mL, 95% CI: -5.7 to -1.3, p=0.008).

When each of the treatment groups was compared to the saline control group, rats in the VAN group had the highest urinary KIM-1, with a significant difference seen after drug dosing on day 1 (Figure 3, 21.5 ng, 95% CI: 3.8 to 39.2, p=0.02), day 2 (29.8 ng, 95% CI: 12.2 to 47.5, p=0.001), day 3 (21.6 ng, 95% CI: 3.9 to 39.2, p=0.02), and day 4 (24.1 ng, 95% CI: 6.4 to 41.7, p=0.008). In a direct comparison to VAN+TZP, rats that received VAN alone had significantly higher urinary KIM-1 after drug dosing on day 1 (18.4 ng, 95% CI: 1.4 to 35.3, p=0.03), day 2 (27.4 ng, 95% CI: 10.4 to 44.3, p=0.002), day 3 (18.8 ng, 95% CI: 1.9 to 35.8, p=0.03), and day 4 (23.2 ng, 95% CI: 6.3 to 40.2, p=0.007). Urinary clusterin was significantly higher among rats in the VAN group on day 2 (2030 ng, 95% CI: 86 to 3974, p=0.04) of the study. Significantly elevated urinary clusterin was also observed among rats in the VAN+TZP group on day 1 (2869 ng, 95% CI: 926 to 4814, p=0.004). Urinary OPN was significantly higher among VAN group rats on day 2 (1.09 ng, 95% CI: 0.33 to 1.86, p=0.005), day 3 (1.35 ng, 95% CI: 0.58 to 2.11, p=0.001), and day 4 (1.73 ng, 95% CI: 0.97 to 2.49, p<0.001). Rats In the VAN+TZP group also had significantly higher urinary OPN on day 3 (0.89 ng, 95% CI: 0.13 to 1.66, p=0.0 2) and day 4 (1.76 ng, 95% CI: 1.0 to 2.53, p<0.001).

**Figure 3:**
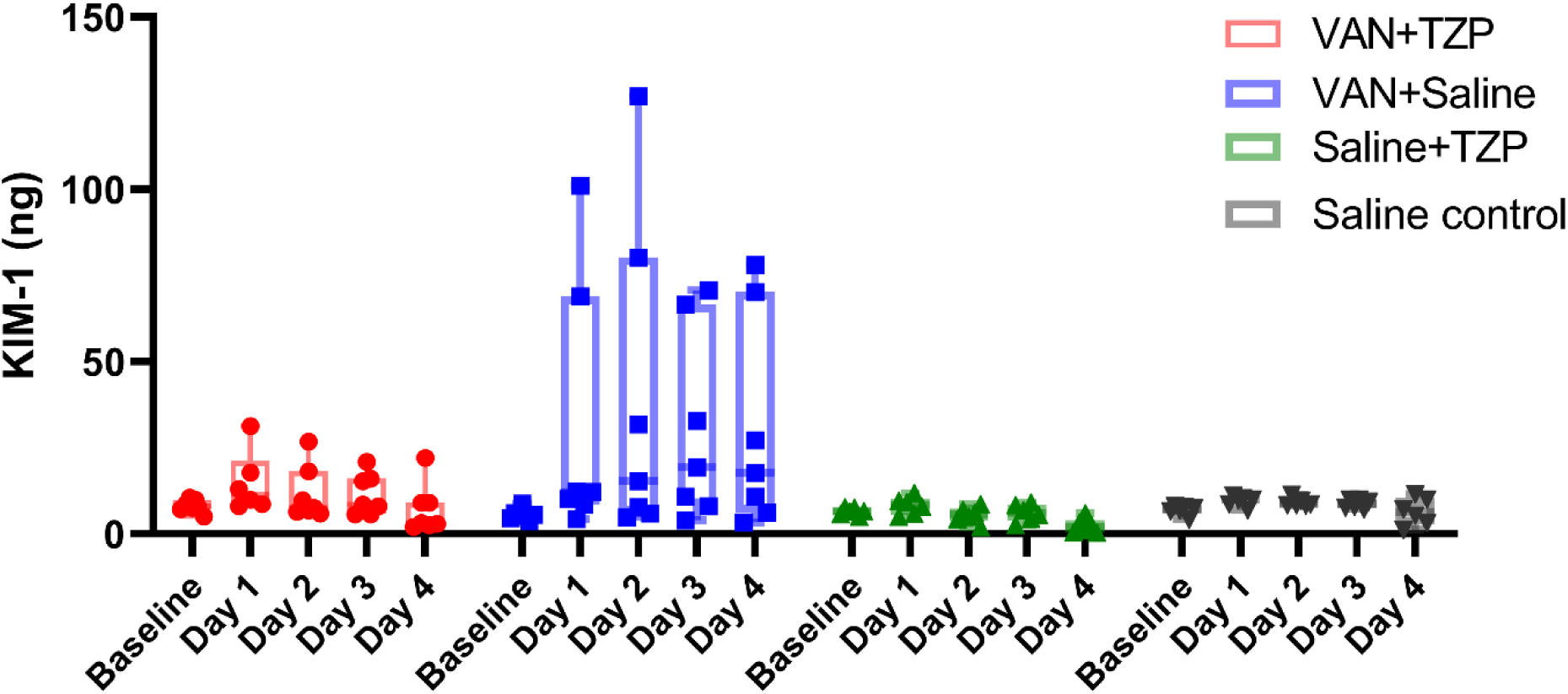
KIM-1 comparison between groups across dosing days. Comparison of individual daily urinary KIM-1 measurements between treatment groups, and across dosing days. Rats in the VAN group had the highest urinary KIM-1, with a significant difference seen after drug dosing on day 1 (21.5 ng, 95% CI: 3.8 to 39.2, p=0.02), day 2 (29.8 ng, 95% CI: 12.2 to 47.5, p=0.001), day 3 (21.6 ng, 95% CI: 3.9 to 39.2, p=0.02), and day 4 (24.1 ng, 95% CI: 6.4 to 41.7, p=0.008). Individual rats are depicted by each data point.

### Correlation between urinary biomarkers of injury and GFR

Spearman’s rank correlation between urinary biomarkers and GFR readings are listed in Table 3. In the VAN group rats, urinary KIM-1 was significantly correlated with decreasing GFR on day 3 (Figure 4, Spearman’s rho: -0.45, p = 0.022) and day 4 (Spearman’s rho: -0.41, p = 0.036). Urinary clusterin was also negatively correlated with GFR on day 3 (Spearman’s rho: -0.42, p=0.033). No significant correlations were observed between urinary OPN and GFR.

**Figure 4:**
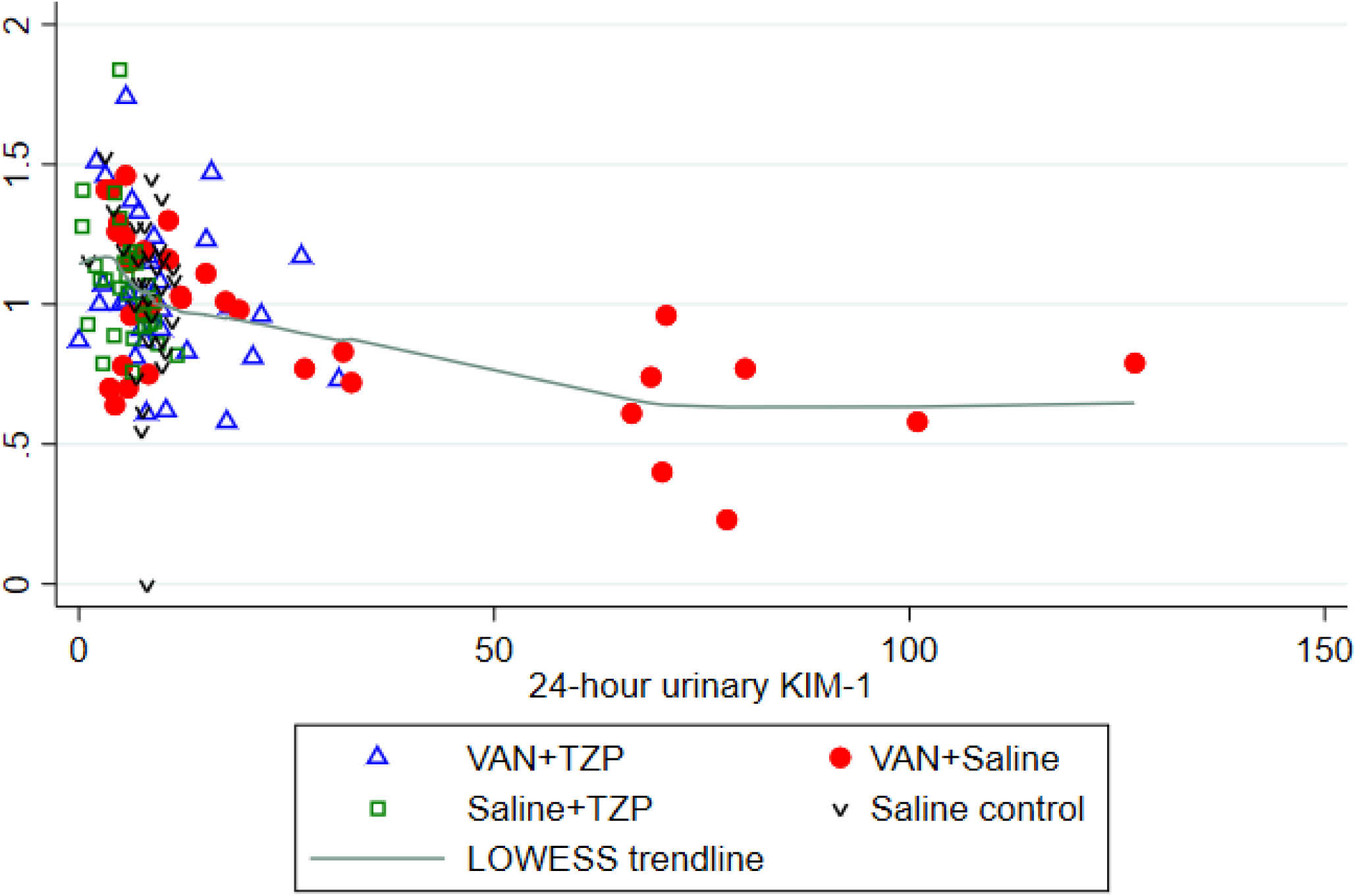
Correlations between urinary KIM-1 and daily GFR. Spearman correlation of GFR with 24-hr urinary KIM-1 levels. Among rats which received VAN+saline, urinary KIM-1 was significantly correlated with decreasing GFR on day 3 (Spearman’s rho: -0.45, p = 0.022) and day 4 (Spearman’s rho: -0.41, p = 0.036). Individual rats are depicted by each data point and the green line depicts the LOWESS trendline.

### Histopathological analysis

Kidney histopathology demonstrated the highest Predictive Safety Testing Consortium (PSTC) scores in the VAN group (median: 2, IQR: 1), followed by the saline control group (median: 1.5, IQR: 1), TZP group (median: 1, IQR: 0), and the VAN+TZP group (median: 1, IQR: 0). The same trend was observed in tubular specific scores with the highest scores observed in the VAN group (median: 2, IQR: 1), followed by the saline control group (median: 1, IQR: 1), TZP group (median: 1, IQR: 0), and the VAN+TZP group (median: 1, IQR: 0). Degenerative renal tubular changes consisting of tubular epithelial necrosis or apoptosis were only observed among rats that received VAN alone. Other changes of tubular epithelial injury, cellular sloughing, and granular intratubular casts, were consistently observed in kidneys from rats that received VAN alone. These indicators of tubular epithelial injury were observed less frequently and at lower severity in kidneys from rats that received VAN+TZP.

## Discussion

In this study, we found that VAN alone led to lower GFRs (as measured by FIT-C sinistrin clearance) after 4 days of drug dosing. Rise of urinary KIM-1, as a marker of proximal tubule injury, occurred as early as day 1 and correlated with the drop in GFR by day 3 among rats that received only VAN. Notably, similar trends in GFR and urinary KIM-1 expression were not observed in rats which received VAN+TZP; in fact, rats which received VAN+TZP recovered both GFR and urinary KIM-1 to baseline levels by day 4 of the experiment. Our results are concordant with previous pre-clinical studies for the injury biomarker KIM-1, in that renal injury was seen earlier and to a greater extent in the VAN alone group compared to the VAN+TZP group (15, 18, 19). Among the assessed urinary biomarkers, KIM-1 was most closely correlated with GFR over the course of the experiment. This is also consistent with our previous findings that higher urinary KIM-1 predicts worse histopathologic damage scores in the rat (15, 17). Clusterin and OPN are more general biomarkers of non-specific tubule or glomerular damage, and both were less correlated with GFR changes in this study (17). These results and their concordance with previous animal studies provide further evidence that VAN+TZP is not associated with additive nephrotoxicity and may even be protective during the initial days of treatment.

To our knowledge, this is the first study to combine a transdermal monitoring device and intravenous fluorescent tracer for measurement of real-time GFR with renal injury biomarkers for vancomycin induced kidney injury in a rat model. Similar to previous studies, we employed our well-established translational rat model to investigate allometrically scaled antibiotic doses and associated toxicity. Utilization of a transdermal monitoring device with an intravenous fluorescent tracer (fluorescein isothiocyanate-labelled sinistrin [FITC-sinistrin]) offers several important advantages over other approaches for estimating kidney function. First, in contrast to timed PKPD studies where sequential serum and/or urine samples must be collected, the transdermal device allows for continuous, non-invasive monitoring of the tracer (which is freely filtered and neither secreted nor reabsorbed by the kidneys in the case of FITC-sinistrin). Second, real-time estimation of GFR allows for the capture of early changes in renal function (i.e. days 1-3). Injury biomarkers are important; however, functional changes are arguably even more relevant for clinical translation. In humans, detectable rises in SCr are seen approximately 2-4 days after the initial renal injury (20, 21). Due to this lag time in SCr rise, early changes in renal function and/or damage may not be detected with conventional measures. Consequently, simultaneous capture of injury biomarkers and unbiased estimates of function allow for maximal translation to clinical utility. It is not expected that real time functional measures of GFR will be available in routine clinical practice in the next few years; hence, understanding the relationship that defines the time course and magnitude of damage by using injury biomarkers will be important for predicting functional changes in clinical practice.

Several considerations for our study should be stated. First because of technical difficulty, one animal did not have adequate GFR data collected on one study day. This animal was still included in our analysis because the statistical methodology that we utilized is flexible and did not require dropping the data (such as with ANOVA methods). The animal provided adequate GFR data on the other experimental days, in addition to serum and terminal kidney samples. No major interpretations change if this animal were in fact excluded from analyses (data not shown). Second, we measured 24-hour urinary volumes, thereby capturing the total amounts of excreted biomarkers (versus obtaining spot concentrations). Findings do not drastically change when analyzed as 24 hour concentrations (Fig 4); however, this is a potential limitation of clinical studies where total urine collections are logistically more difficult and less common. Future studies to enhance clinical translation may benefit from employing blood biomarkers where dilution and sample collection labor is less of a factor, compared to urinary biomarkers. Despite these concerns, it is notable that KIM-1 in the rat is a homologue of KIM-1b in humans. Human studies have corrected for urinary volume by standardizing to urinary creatinine (22); however that is less necessary in well controlled animal studies where dilution can be fully calculated. Finally, while urinary KIM-1 in the rat has been linked to PK/PD predictions in humans, it is less clear what changes in rat GFR mean (23). While there is not a direct numeric translation for GFR, this study demonstrates proof of principle that GFR changes exist with VAN but do not with VAN +TZP at the administered doses. Minimally, one can conclude that TZP does not worsen kidney function or injury when added to VAN in the rat. Ultimately, these results may have clinical implications beyond the specific combination of VAN+TZP since specific renal injury biomarkers may be employed in monitoring other nephrotoxic medications. Further clarification of urinary biomarkers of renal injury is needed to elucidate specific biomarker elevations and expression patterns associated with various nephrotoxins, prior to realized functional kidney changes.

In our translational rat model assessing kidney function, rats that received VAN had a significant decline in GFR by day 4 whereas no decline in kidney function was observed in other groups (including VAN+TZP). In conclusion, the addition of TZP to VAN does not worsen kidney function, as measured by GFR. These data are consistent with animal injury models and demonstrate that VAN+TZP is not associated with decreased glomerular function nor additive kidney injury, when compared to VAN. Clinical studies that wish to assess the impact of TZP added to VAN should directly measure renal injury biomarkers and estimate GFR in an unbiased methodology that does not employ serum creatinine.

## Materials and Methods

### Experimental design and animals

The experimental methods were similar to those we have previously reported for our employed translational rat model (15, 18, 19). All experiments were conducted at Midwestern University in Downers Grove, Illinois, in compliance with the National Institutes of Health Guide for the Care and Use of Laboratory Animals (24) and were approved under the Institutional Animal Care and Use Committee protocol #3151. In brief, male Sprague-Dawley rats (n=26; age: approximately 8 to 10 weeks; mean weight: 274.8 g) were housed in a light- and temperature-controlled room(15, 18, 19) for the duration of the study and allowed free access to water and food (Supplemental figure 1). All animals were placed in metabolic cages (Nalgene, Rochester, NY, USA) for 24-hour urine collection, starting prior to dosing (day 0) and sampled every day for a period of 4 days. Animals were assigned to one of four treatment groups in which they received either VAN 150 mg/kg/day intravenously, over 2 minutes (n=7), TZP 1400mg/kg/day intraperitoneal (n=6), VAN+TZP (n=7), or saline (n=6) for 4 days (Supplemental figure 2). At baseline (prior to drug dosing) and following drug dosing on each day, FITC-sinistrin 5mg/100g was administered intravenously to quantify GFR as described below. Following completion of the dosing protocol, rats were sacrificed and underwent nephrectomies.

### Chemicals and reagents

Rats were administered clinical grade vancomycin (Lot number: 167973; Fresenius Kabi, Lake Zurich, IL, USA), piperacillin-tazobactam (Lot number: 1PU19022; Apollo, Palm Beach Gardens, USA), and normal saline for injection (Hospira, Lake Forest, IL, USA). VAN and TZP were prepared by weighing and dissolving the powder in normal saline to achieve final concentrations of 100 mg/mL and 500 mg/mL, respectively. FITC-sinistrin was prepared by weighing and dissolving the powder in sterile 1x phosphate buffered saline to achieve a final concentration of 40 mg/mL.

### Blood, urine, and kidney sampling

Double jugular vein catheters were surgically implanted 72 h prior to protocol initiation. One catheter was dedicated to blood sample draws, while drug dosing occurred via the other catheter. Blood samples were obtained from the catheter at pre-specified timepoints (0 and 240 mins). Each sample (0.2 mL/aliquot) was replaced with an equivalent volume of normal saline (NS) for maintenance of euvolemia. Blood samples were prepared as plasma with disodium ethylenediaminetetraacetic acid (EDTA) salt dihydrate (Lot: 19C1856562; Sigma-Aldrich Chemical Company, Milwaukee WI, USA) and centrifuged at 3000 g for 10 minutes (Thermo Fisher Scientific, Waltham, MA, USA). Supernatants were collected and frozen at -80°Cfor batch analysis.

Urine samples were collected and volume was measured every 24 hours, starting from day 0. Urine samples were centrifuged at 400 g for 5 minutes, and the resulting supernatant was collected, aliquoted, and stored at - 80°Cuntil batch analysis of renal biomarkers.

Rats underwent terminal nephrectomies once the dosing protocol was completed (Supplemental figure 3). Left kidneys were formalin fixed for batch histopathological analysis and right kidneys were flash frozen in liquid nitrogen and stored at -80°C.

### GFR measurement

A transdermal monitoring device and intravenous FITC-sinistrin was used to obtain real-time GFR readings. This methodology is an improvement over traditional markers of kidney function such as SCr and urine output. FITC-sinistrin is a biomarker for kidney function that shows comparable kinetics to gold-standard markers such as inulin, with superior handling and administration characteristics (25). In addition, the ability to estimate real-time GFR allows for the capture of much earlier changes in kidney function compared to creatinine (i.e. days 1-3). Transdermal sensors (MediBeacon GmBH, Mannheim, Germany) and FITC-sinistrin (Lot number: VE17200811; Fresenius Kabi, Hamburg, Germany) were used for GFR estimation. A small area on the dorsal side of each rat was depilated (Nair; Church & Dwight Co, Ewing, NJ, USA) on day 0, 24 hours prior to the start of the experimental protocol. FITC-sinistrin 5mg/100g was administered daily as an intravenous push. Fluorescence was monitored continuously (one measurement every 8 seconds) via the transdermal sensor for 2 hours at baseline on day 1, and 4 hours after each daily dose (26). At the completion of the measurement period, the sensor was removed, and data was transferred to a computer for analysis and storage. Data were analyzed using the MB Studio software (MediBeacon GmBH, Mannheim, Germany). A 3-compartment model with linear baseline correction was fit to determine FITC-sinistrin clearance vis-à-vis GFR.

### Determination of urinary biomarkers of AKI

Urine samples were batch analyzed to determine urinary concentrations of KIM-1, clusterin, and OPN. Microsphere-based Luminex xMAP technology was used for the determination of urinary protein biomarkers as previously described (17, 27). Urine samples were allowed to thaw at ambient room temperature, aliquoted into 96-well plates, and mixed with Milliplex MAP rat kidney toxicity magnetic bead panel 1 (EMD Millipore Corporation, Charles, MO, USA). A separate standard curve was prepared and run with each assay plate, per the manufacturer’s instructions. Results were analyzed and urinary biomarker concentrations were determined using the manufacturer’s software which utilizes flexible five-parameter (linear and logarithmic scale) curve-fitting models (Milliplex Analyst 5.1; VigeneTech, Carlisle, MA, USA).

### Histopathological analysis of kidneys

Formalin-fixed kidneys were sent for histopathological analysis (IDEXX Bioanalytics, Columbia, MO, USA). Blinded samples were prepared and scored according to the PSTC semi-quantitative grading system. Scores range from 0 to 5, with 0 indicating no abnormality, 1 indicating minimal/very few/very small abnormalities, 2 indicating mild/slight/few/small abnormalities, 3 indicating moderate/moderate/number/moderate size abnormalities, 4 indicating marked/many/large abnormalities, and 5 indicating massive/extended number/extended size abnormalities noted. The worst overall score, representing total gross kidney damage, and worst tubular scores were utilized in our analysis.

### Statistical analysis

A mixed-effects, restricted maximum likelihood estimation regression was used to compare urine output, mean weight loss, GFR, and urinary biomarkers among the treatment groups, with repeated measures occurring over days; measures were repeated at the level of the individual rat (Stata version 16.1, StataCorp LLC, College Station, TX, USA). Spearman’s rank correlation coefficient with a Bonferroni correction was used to assess correlations between kidney injury (e.g. KIM-1, clusterin, OPN) and function (e.g. GFR) by treatment day. All tests conducted were two-tailed, with an *a priori* level of statistical significance set at α = 0.05.

## Acknowledgements

The research reported in this publication was supported in part by the National Institute of Allergy and Infectious Diseases under award number R21-AI149026 (authors MS, GP). The content is solely the responsibility of the authors and does not necessarily represent the official views of the National Institutes of Health.

MS reports ongoing research contracts with Nevakar and SuperTrans Medical as well as having filed patent US10688195B2. All other authors have no other related conflicts of interest to declare.

## Supplemental figures

**Supplemental figure 1:**
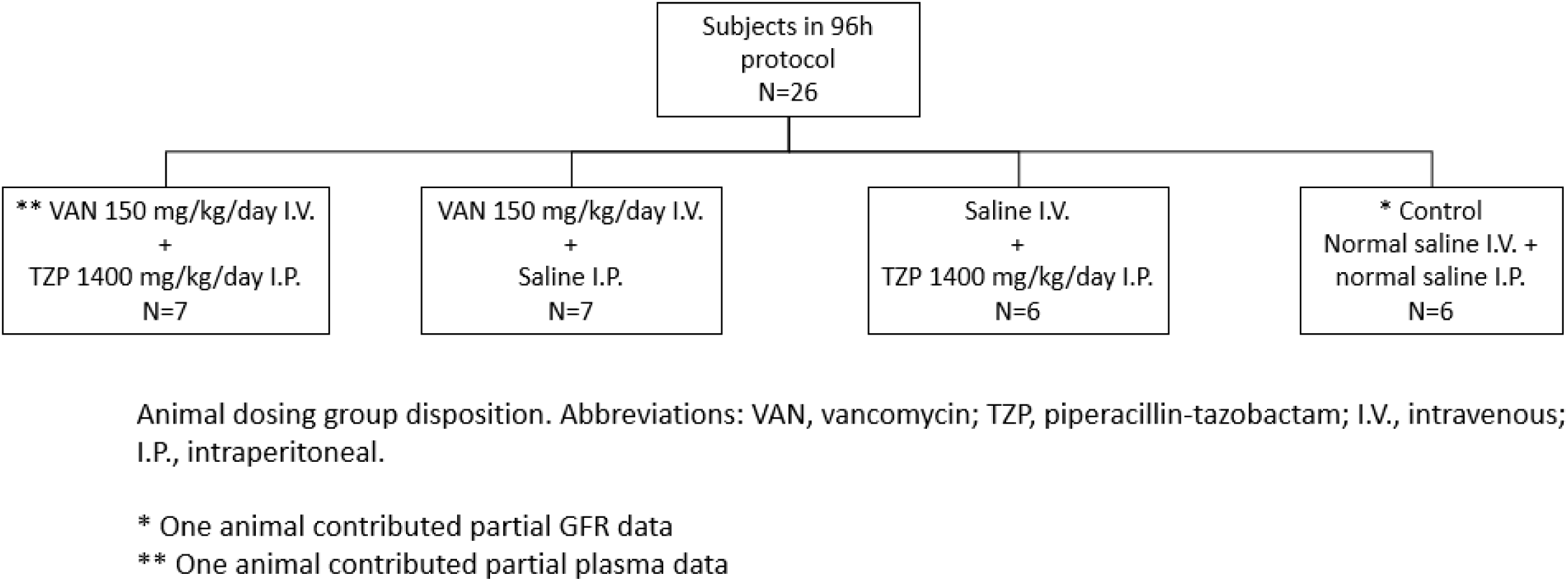
Study treatment groups assignment

**Supplemental figure 2:**
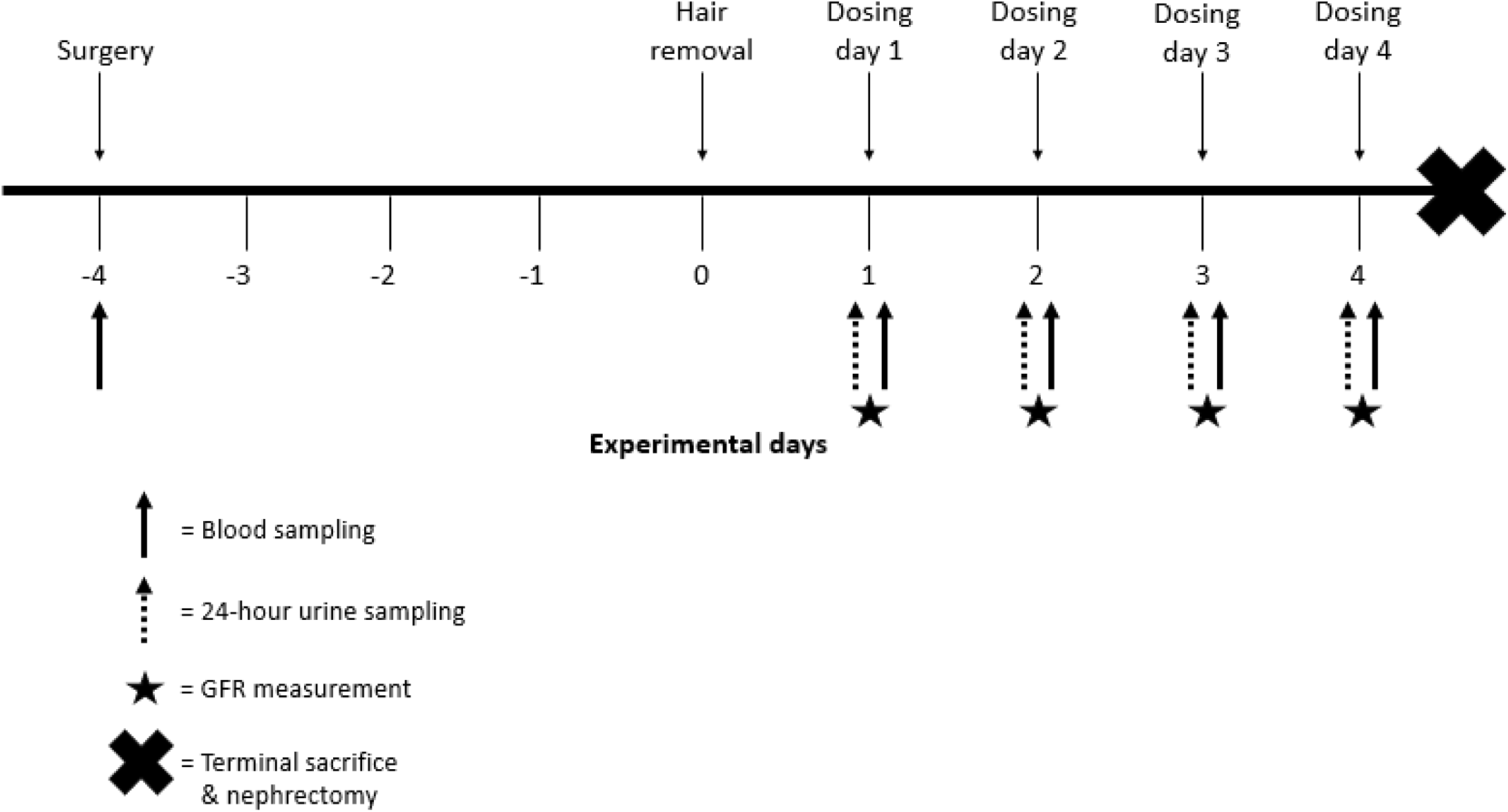
Overall experimental timeline

**Supplemental figure 3:**
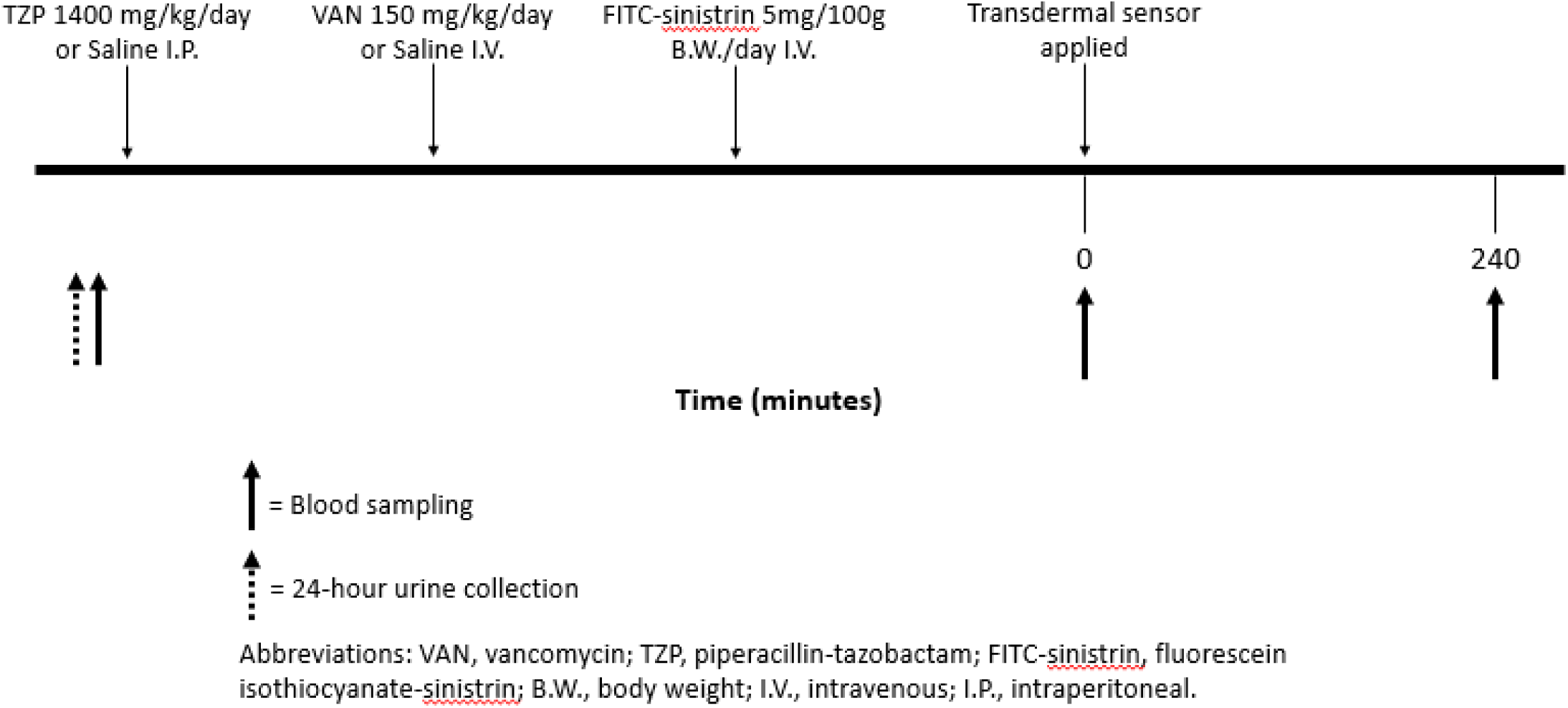
Daily experimental timeline

**Supplemental figure 4:**
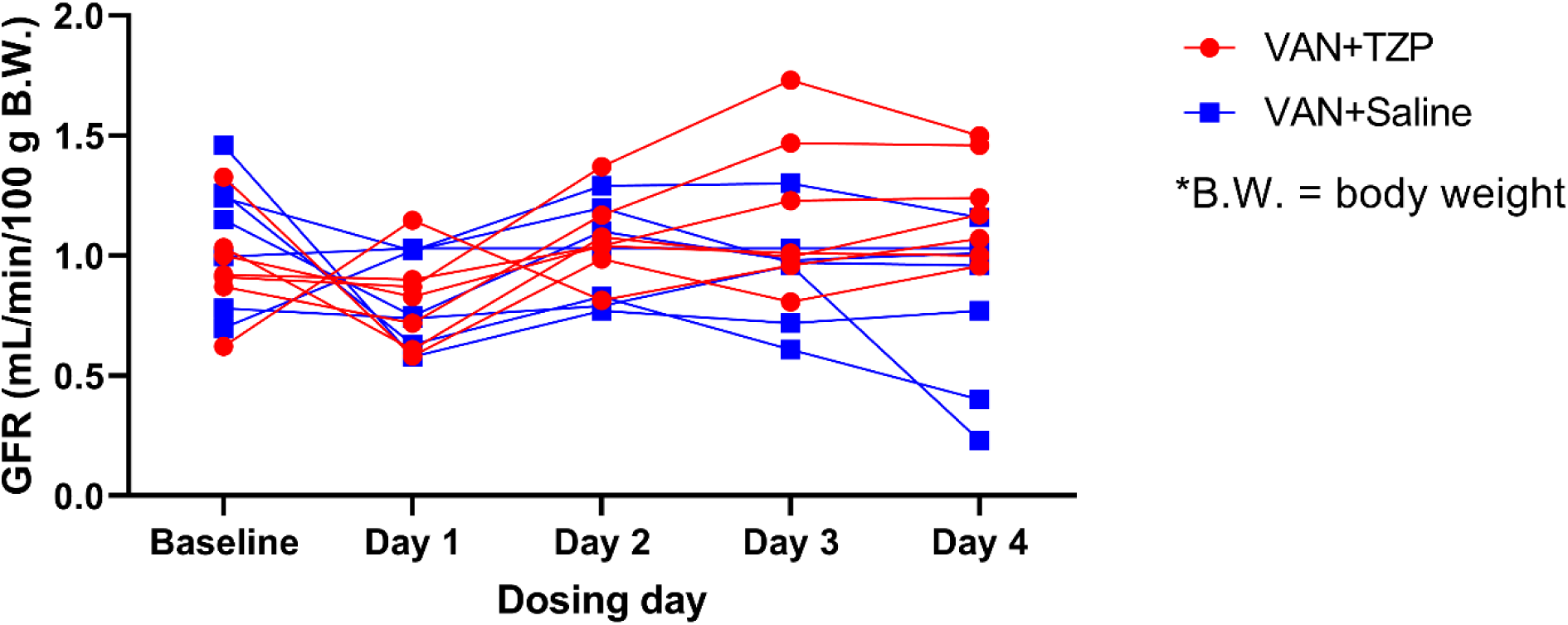
GFR comparison between VAN+TZP and VAN+Saline groups. Direct comparison of individual GFRs in the VAN+TZP and VAN alone groups. Only rats which received VAN had a significant decline in GFR by day 4 (−0.35 mL/min/100 g body weight, 95% CI: - 0.63 to -0.07, p=0.013).

**Supplemental figure 5:**
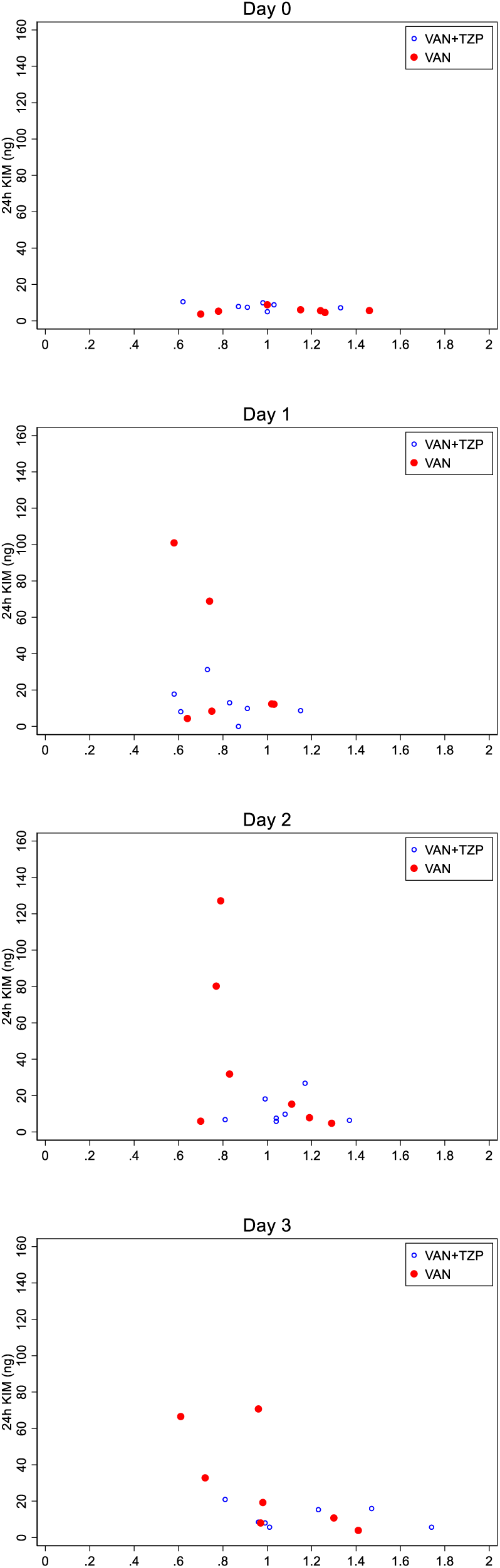
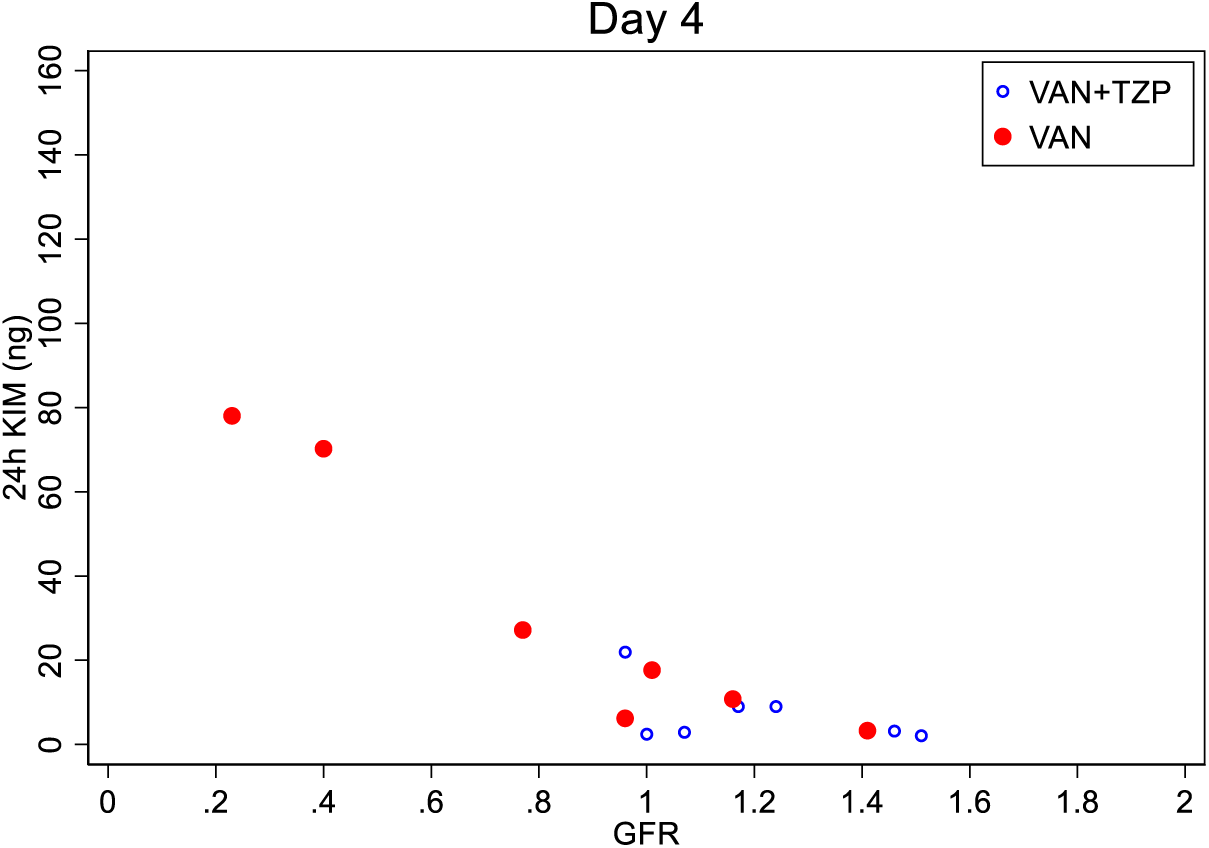
Spearman correlations between GFR and 24-hour urinary KIM-1 across treatment days for VAN+TZP and VAN alone groups. On day 0 (baseline), all rats in both VAN+TZP and VAN groups had similarly low KIM-1 levels prior to administration of study drug. After drug dosing on day 1 and day 2, we begin to see VAN group rats (red data points) with decreased GFR and higher KIM-1 levels. Following study drug dosing on day 3, urinary KIM-1 for the VAN group was significantly correlated with decreasing GFR (Spearman’s rho: -0.45, p = 0.022). A similar correlation was observed between GFR and KIM-1 for the VAN group rats on day 4 (Spearman’s rho: -0.41, p = 0.036).

**Supplementary Table 1:** Summary of weight loss, urinary output, and urinary biomarkers.

| Characteristic | Treatment group |  |  |  |
| --- | --- | --- | --- | --- |
|  | VAN | TZP | VAN+TZP | Saline control |
| Number of animals | 7 | 6 | 7 | 6 |
| Baseline weight (g), mean ± SD | 274.04 ± 8.46 | 280.53 ± 13.29 | 277.46 ± 5.19 | 278.25 ± 11.04 |
| Weight change (g), mean ± SD | -1.39 ± 6.09 | -2.27 ± 6.25 | -1.59 ± 5.44 | +5.50 ± 9.06 |
| Urine output (mL), mean ± SD | | | | |
| Baseline | 8.73 ± 3.09 | 10.95 ± 4.78 | 11.43 ± 3.95 | 8.1 ± 1.42 |
| Day 1 | 15.67 ± 3.7 | 12.98 ± 2.88 | 17.1 ± 3.07 | 11.58 ± 1.59 |
| Day 2 | 8.83 ± 2.63 | 6.40 ± 2.54 | 9.13 ± 2.6 | 8.38 ± 2.18 |
| Day 3 | 7.64 ± 3.22 | 7.73 ± 3.53 | 10.34 ± 3.57 | 7.15 ± 1.37 |
| Day 4 | 13.4 ± 8.95 | 4.43 ± 3.65 | 11.3 ± 5.53 | 4.49 ± 3.27 |
| KIM-1 (ng), median (IQR) |  |  |  |  |
| Baseline | 5.58 (4.59 to 6.09) | 6.43 (5.7 to 7.65) | 7.91 (7.2 to 9.94) | 7.22 (6.73 to 7.92) |
| Day 1 | 12.2 (8.37 to 68.8) | 8.44 (5.81 to 9.46) | 9.88 (8.12 to 17.8) | 9.97 (8.74 to 10.44) |
| Day 2 | 15.3 (5.91 to 80.2) | 5.53 (4.18 to 6.88) | 7.59 (6.4 to 18.2) | 8.58 (8.4 to 9.70) |
| Day 3 | 19.3 (8.0 to 66.6) | 6.11 (4.26 to 8.31) | 8.5 (5.7 to 15.9) | 8.62 (7.69 to 10.1) |
| Day 4 | 17.7 (6.21 to 70.2) | 1.92 (0.4 to 3.16) | 3.2 (2.46 to 9.03) | 6.3 (3.2 to 9.98) |
| Clusterin (ng), median (IQR) |  |  |  |  |
| Baseline | 2282 (1466 to 3004) | 2734 (2120 to 3080) | 3750 (2608 to 5164) | 1743 (1074 to 2563) |
| Day 1 | 4362 (2021 to 4505) | 2314 (1962 to 2829) | 3879 (2517 to 8381) | 2186 (1281 to 2496) |
| Day 2 | 2564 (1431 to 7520) | 1168 (787 to 1401) | 2522 (1291 to 3150) | 1380 (1075 to 2342) |
| Day 3 | 2139 (1530 to 5572) | 1268 (1139 to 1421) | 2534 (1497 to 4592) | 1160 (832 to 1456) |
| Day 4 | 1199 (833 to 2429) | 219 (129 to 1895) | 746 (573 to 3269) | 616 (401 to 1376) |

| Osteopontin (ng), median (IQR) |  |  |  |  |
| --- | --- | --- | --- | --- |
| Baseline | 0.31 (0 to 0.68) | 0.38 (0.15 to 0.60) | 0.38 (0 to 0.837) | 0.29 (0.21 to 0.31) |
| Day 1 | 0.99 (0.27 to 1.32) | 0.86 (0.48 to 1.35) | 0.58 (0.45 to 1.68) | 0.33 (0.22 to 0.48) |
| Day 2 | 1.33 (0.84 to 2.58) | 0.77 (0.56 to 1.12) | 0.88 (0.65 to 1.40) | 0.62 (0.44 to 0.70) |
| Day 3 | 2.16 (0.67 to 2.34) | 1.01 (0.42 to 1.46) | 1.12 (1.09 to 1.68) | 0.37 (0.17 to 0.43) |
| Day 4 | 0.54 (0.23 to 1.02) | 0.23 (0.06 to 0.50) | 0.49 (0.36 to 0.81) | 0.15 (0.11 to 0.19) |
| GFR (mL/min/100 g B.W.), mean ± SD | | | | |
| Baseline | 1.08 ± 0.27 | 1.09 ± 0.38 | 0.96 ± 0.21 | 0.91 ± 0.53 |
| Day 1 | 0.79 ± 0.19 | 0.98 ± 0.18 | 0.81 ± 0.19 | 0.97 ± 0.26 |
| Day 2 | 0.95 ± 0.24 | 1.06 ± 0.12 | 1.07 ± 0.17 | 1.06 ± 0.13 |
| Day 3 | 0.99 ± 0.29 | 1.15 ± 0.14 | 1.17 ± 0.33 | 1.01 ± 0.22 |
| Day 4 | 0.85 ± 0.42 | 1.11 ± 0.23 | 1.20 ± 0.22 | 1.24 ± 0.18 |

